# Fine-scale human population structure in southern Africa reflects ecological boundaries

**DOI:** 10.1101/038729

**Authors:** Caitlin Uren, Minju Kim, Alicia R Martin, Dean Bobo, Christopher R Gignoux, Paul D van Helden, Marlo Möller, Eileen G Hoal, Brenna M Henn

## Abstract

Recent genetic studies have established that the KhoeSan populations of southern Africa are distinct from all other African populations and have remained largely isolated during human prehistory until about 2,000 years ago. Dozens of different KhoeSan groups exist, belonging to three different language families, but very little is known about population history within southern Africa. We examine new genome-wide polymorphism data and whole mitochondrial genomes for more than one hundred South Africans from the ≠Khomani San and Nama populations of the Northern Cape, analyzed in conjunction with 19 additional southern African populations. Our analyses reveal fine-scale population structure in and around the Kalahari Desert. Surprisingly, this structure does not always correspond to linguistic or subsistence categories as previously suggested, but rather reflects the role of geographic barriers and the ecology of the greater Kalahari Basin. Regardless of subsistence strategy, the indigenous Khoe-speaking Nama pastoralists and the N|u-speaking ≠Khomani (formerly hunter-gatherers) share recent ancestry with other Khoe-speaking forager populations that forms a rim around the Kalahari Desert. We reconstruct earlier migration patterns and estimate that the southern Kalahari populations were among the last to experience gene flow from Bantu-speakers, approximately 14 generations ago. We conclude that local adoption of pastoralism, at least by the Nama, appears to have been primarily a cultural process with limited impact from eastern African genetic diffusion.

## Introduction

The indigenous populations of southern Africa, referred to by the compound ethnicity “KhoeSan” (Schlebusch 2010), have received intense scientific interest. This interest is due both to the practice of hunter-gatherer subsistence among many groups - historically and to present-day - and recent genetic evidence suggesting that the ancestors of the KhoeSan diverged early on from other African populations (Behar *et al*. 2008; Tishkoff *et al*. 2009; Henn *et al*. 2011, 2012; Pickrell *et al*. 2012; Veeramah *et al*. 2012; Barbieri *et al*. 2013). Genetic data from KhoeSan groups has been extremely limited until very recently, and the primary focus has been on reconstructing early population divergence. Demographic events during the Holocene and the ancestry of the Khoekhoe-speaking pastoralists have received limited, mostly descriptive, attention in human evolutionary genetics. However, inference of past population history depends strongly on understanding recent population events and cultural transitions.

The KhoeSan comprise a widely distributed set of populations throughout southern Africa speaking, at least historically, languages from one of three different linguistic families - all of which contain click consonants rarely found elsewhere. New genetic data indicates that there is deep population divergence even among KhoeSan groups (Pickrell *et al*. 2012; Schlebusch *et al*. 2012, 2013; Schlebusch and Soodyall 2012; Barbieri *et al*. 2013), with populations living in the northern Kalahari estimated to have split from southern groups 30,000-35,000 years ago (Pickrell *et al*. 2012; Schlebusch *et al*. 2012; Schlebusch and Soodyall 2012). Pickrell et al. (2012) estimates a time of divergence between the northwestern Kalahari and southeastern Kalahari population dating back to 30,000 years ago; “northwestern” refers to Juu-speaking groups like the !Xun and Ju/’hoansi while “southeastern” refers to Taa-speakers. In parallel, Schlebusch et al. (2012) also estimated an ancient time of divergence among the KhoeSan (dating back to 35,000 ya), but here the southern groups include the ≠Khomani, Nama, Karretjie (multiple language families) and the northern populations refer again to the !Xun and Ju/’hoansi. Thus, KhoeSan populations are not only strikingly isolated from other African populations but they appear geographically structured among themselves. To contrast this with Europeans, the ≠Khomani and the Ju/’hoansi may have diverged over 30,000 *ya* but live only 1,000 km apart, roughly the equivalent distance between Switzerland and Denmark whose populations have little genetic divergence (Novembre et al. 2008). However, it is unclear how this ancient southern African divergence maps on to current linguistic and subsistence differences among populations, which may have emerged during the Holocene.

In particular, the genetic ancestry of the Khoe-speaking populations and specifically the Khoekhoe, (e.g. Nama) who practice sheep, goat and cattle pastoralism, remains a major open question. Archaeological data has been convened to argue for a demic migration of the Khoe from eastern African into southern Africa, but others have also argued that pastoralism represents cultural diffusion without significant population movement (Boonzaier 1996; K. C. MacDonald 2000; Robbins *et al*. 2005; Sadr 2008, 2015; Dunne *et al*. 2012; Pleurdeau *et al*. 2012; Jerardino *et al*. 2014). Work by Breton et al. and Macholdt et al. discovered the presence of lactase persistence alleles in KhoeSan groups, especially frequent in the Nama (20%), which clearly derive from eastern African pastoralist populations (Breton *et al*. 2014; Macholdt *et al*. 2014). This observation, in conjunction with other Y-chromosome and autosomal data (Henn *et al*. 2008), were used to argue that pastoralism in southern Africa was another classic example of demic diffusion. However, previous work is problematic in that it tended to focus on single loci [MCM6/LCT, Y-chromosome] subject to drift or selection, overall eastern African autosomal ancestry remains minimal and the distribution of ancestry informative markers is dispersed between both pastoralist and hunter-gatherer populations. Here, we present a comprehensive study of recent population structure in southern Africa and clarify fine-scale structure beyond “northern” and “southern” geographic descriptors. We then specifically test whether the Khoe-speaking Nama pastoralists derive their ancestry from eastern Africa, the northeastern Kalahari Basin, or far southern Africa. Our results suggest that the ecology of southern Africa is a better explanatory feature than either language, clinal geography or subsistence on its own.

## Results

To resolve fine-scale population structure and migration events in southern Africa, we generated genome-wide data from three South African populations. We genotyped ≠Khomani San (*n*=75), Nama (*n*=13) and South African Coloured (SAC) (*n*=25) individuals on the Illumina OmniExpress and OmniExpressPlus SNP array platforms. Sampling locations are listed in Table S1. These data were merged with HapMap3 (joint Illumina Human1M and Affymetrix SNP 6.0) (Consortium 2010), HGDP (Illumina 650Y) data (Li *et al*. 2008) and the dataset from Petersen et al. (Petersen *et al*. 2013) (Illumina HumanOmni1-Quad Beadchips), resulting in an intersection of ∽320k SNPs for 852 individuals from 21 populations. In addition, we used the Affymetrix Human Origins SNP Array generated as part of Pickrell et al. (Pickrell *et al*. 2012) and Lazaridis et al. (Lazaridis *et al*. 2014), including **n*=9* ≠Khomani San individuals from our collection and encompassing over 396 individuals from 33 populations. Whole mitochondrial genomes were generated from offtarget reads from exome-and Y chromosome-capture targeted short read sequencing. Reads were mapped to GRCh37 which uses the revised Cambridge reference sequence (rCRS). Only individuals with greater than 7x haploid coverage were included in the analysis: ≠Khomani San (*n*=64) and Nama (*n*=31). In this study, we address population structure among southern African KhoeSan, the genetic affinity of the Khoe, and when and how pastoralism diffused into southern Africa.

### Population Structure in Southern African KhoeSan Populations

Unsupervised population structure analysis identifies 5 distinct, spatially organized ancestries among the sampled twenty-one southern African populations. These ancestries were inferred from the Affymetrix Human Origins dataset using ADMIXTURE (Figure S1) (Alexander *et al*. 2009). Multi-modality per *k* value was assessed using *pong* (Behr *et al*. 2015) and results from k=10 are discussed below (6/10 runs assigned to the major mode, 3/10 other runs involved cluster switching only within East Africa). Visualization of these ancestries according to geographic sampling location specifically demonstrates fine-scale structure in and around the Kalahari Desert **(Figure 1).** While prior studies have argued for a northern versus southern divergence of KhoeSan populations (Pickrell *et al*. 2012; Schlebusch *et al*. 2012; Schlebusch and Soodyall 2012; Barbieri *et al*. 2013, 2014), the structure inferred from our dataset indicates a more complex pattern of divergence and gene flow. Even recent migration events into southern Africa remain structured, consistent with ecological boundaries to gene flow. The distribution of the five ancestries corresponds to: a northern Kalahari ancestry, central Kalahari ancestry, circum-Kalahari ancestry, a northwestern Namibian savannah ancestry and ancestry from eastern Bantu-speakers **(Figure 1**). This geographic patterning does not neatly correspond to linguistic or subsistence categories, in contrast to previous discussions (Pickrell *et al*. 2012; Schlebusch *et al*. 2012; Barbieri *et al*. 2014).

**Figure 1:**
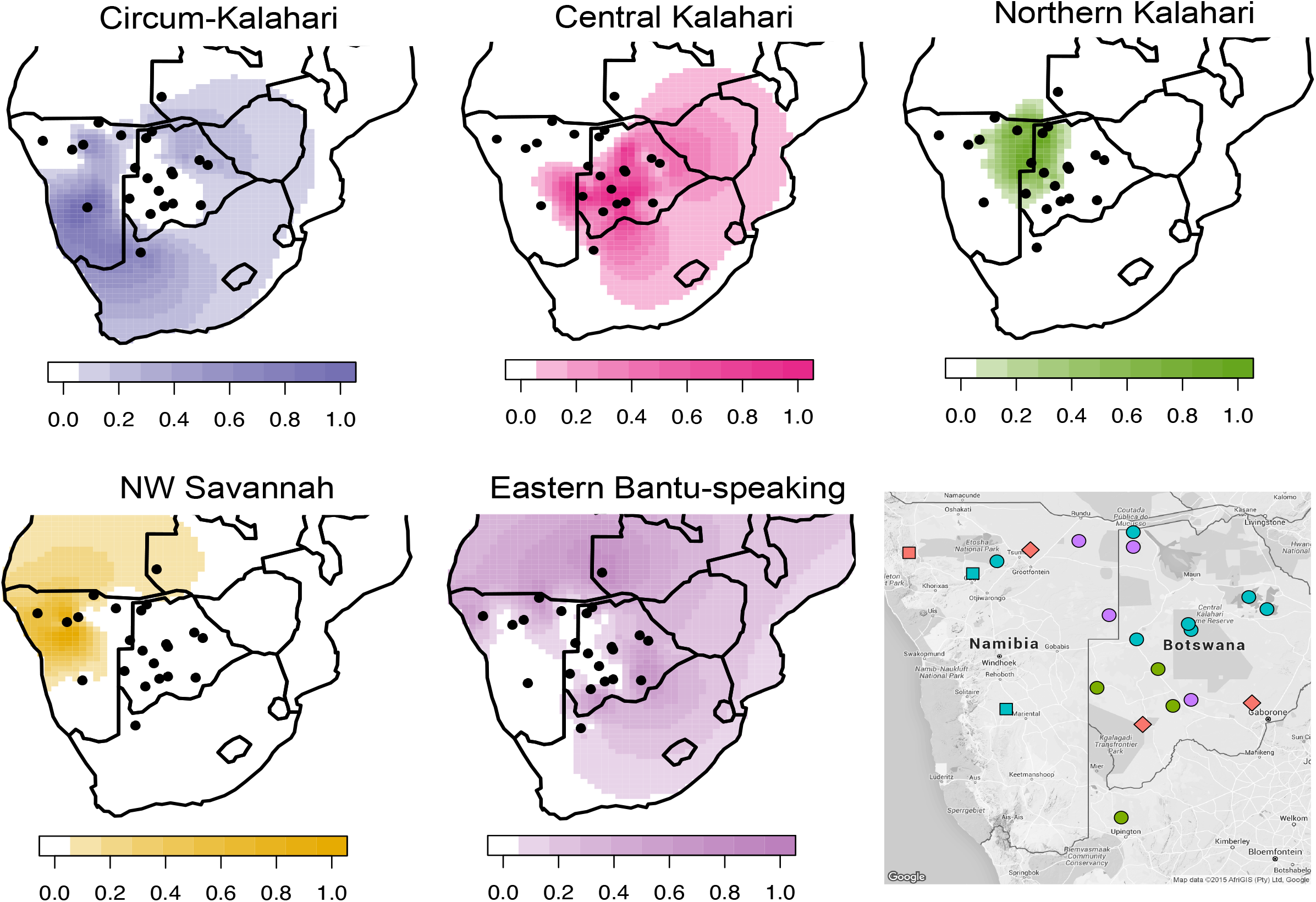
Five spatially distinct ancestries indicate deep population structure in southern Africa. Surface plots of the ancestry proportions from the Affymetrix Human Origins dataset were interpolated across Africa. The X and Y axes are latitude and longitude, respectively. Black dots represent the sampling location of populations in southern Africa. The 3^rd^ dimension in each map [depth of color] represents the mean ancestry proportion for each group calculated from ADMIXTURE *k* = 10 using unrelated individuals, and indicated in the color keys as 0% to 100% for five specific *k* ancestries. The topographic map indicates the subsistence strategy and language of each population. Colors represent language families: green= Tuu speakers, red= Niger-Congo speakers, blue= Khoe speakers and purple= Kx’a speakers. Shapes represent subsistence strategies: circle= hunter-gatherers, square= pastoralists and diamond= agropastoralists.

The northern Kalahari ancestry is the most defined of these ancestries, encompassing several forager populations such as the Ju/’hoansi, !Xun, Khwe, Naro and to a lesser extent the Khoekhoe-speaking Hai||om. While these populations are among the best-studied KhoeSan in anthropological texts with particular reference to cultural similarities (Dornan 1925; Bleek 1928; Schapera 1934; Barnard 1992a), they represent only a fraction of the diversity among Khoisan-speaking populations. We note that this cluster includes Kx’a (Juu), Khoe-Kwadi and Khoekhoe speakers, suggesting that language interacts in a complex fashion with other factors such as subsistence strategy and ecology. The Hai||om are thought to have shifted to speaking Khoekhoe from an ancestral Juu-based language. The central Kalahari ancestry occupies a larger geographical area throughout the Kalahari basin, with its highest frequency among the Taa-speakers: G|ui, G||ana, ≠Hoan and Naro. This ancestry spans all three Khoisan language families (Table S1), at considerable frequency in each, yet all are primarily foragers.

The third ancestry cluster is represented by southern KhoeSan populations distributed along the rim of the Kalahari Desert (**Figure 1**) – referred to here as the “circum-Kalahari ancestry”. The circum-Kalahari ancestry is at its highest frequency in the Nama and ≠Khomani, and significant representation in the Hai||om, Khwe, !Xun and Shua. This ancestry spans all linguistic and subsistence strategies. We propose that the circum-Kalahari is better explained by ecology than alternative factors such as language or recent migration. Specifically, we find the Kalahari Desert is an ecological boundary to gene flow, thus gene flow occurred more frequently among populations on the rim - potentially as the result of larger effective population sizes. The circum-Kalahari ancestry is not easily explained by a pastoralist Khoekhoe dispersal. This spatially distinct ancestry is common in both forager and pastoralist groups, indeed all of the circum-Kalahari populations were historically foragers (except for the Nama). Therefore, to support a Khoekhoe dispersal model, we would have to posit an adoption of pastoralism by a northeastern group, leading to demic expansion around the Kalahari, with subsequent reversion to foraging the the majority of the circum-Kalahari groups; this scenario seems unlikely.

Finally, our analysis reveals two additional ancestries outside of the greater Kalahari Basin: one ancestry composed of Bantu-speakers, frequent to the north, east and southeast of the Kalahari; and a second composed of Himba, Ovambo, and Damara ancestry in northwestern Namibia distributed throughout the mopane savannah. Interestingly, the Damara are a Khoekhoe-speaking population of former foragers (later in servitude to the Nama pastoralists) whose ancestry is unclear *(see below)*.

We used our data and the Affymetrix HumanOrigins dataset containing the greatest number of KhoeSan populations to date, to test whether language or geography better explains genetic distance. The genetic data were compared to a phonemic distance matrix (Jaccard 1908) as well as geographic distances between each population (Table S3). In order to test whether genetic distance (F_st_) was associated with geography or language, we performed a partial Mantel test for the relationship between F_st_ and language (Creanza *et al*. 2015) accounting for geographic distance among 11 KhoeSan populations. This result was not significant *(r*=0.06, *p*=0.30). Although an association between F_st_ and geographical distances within Africa has been documented (Ramachandran *et al*. 2005; Tishkoff *et al*. 2009; Creanza *et al*. 2015), a Mantel test for the relationship between F_st_ and pairwise geographic distance in our dataset was also null (*r*=0.021, *p*=0.38) reflecting the non-linear aspect of shared ancestry in southern Africa as seen in **Figure 1.**

### A Divergent Southern KhoeSan Ancestry

Spatially distinct ancestries are also supported by principal components analysis (PCA) **(Figure 2 and S2).** As observed previously in a global sample of populations (Henn *et al*. 2011), the KhoeSan anchor one end of PC1 opposite to Eurasians. PC2 separates other African populations from the KhoeSan, including western Africans as well as central and eastern African hunter-gatherers. PC3 separates the Ju/’hoansi and !Xun (northern Kalahari) from ≠Hoan, Taa-speakers and Khoe-speakers, with other KhoeSan populations intermediate. This separation of northern (Ju/’hoansi) and southern (Taa and Khoe speakers) KhoeSan populations has been observed by Schlebusch et al. (Schlebusch *et al*. 2012) and Pickrell et al. (Pickrell *et al*. 2012). We estimate that this trans-Kalahari genetic differentiation from the inferred ancestral allele frequencies (Figure S3) is substantial (F_ST_ = 0.05) concordant with prior estimates of a 20,000-35,000 year divergence between the ancestries, far greater than that seen across Europe (Novembre *et al*. 2008). We verify this divergence between the northern Kx’a-speakers and the shared Nama and ≠Khomani ancestry in a new, second sample of Nama, from South Africa rather than Namibia (Table S1, Figure S3). This southern KhoeSan ancestry is also present in admixed Bantu-speaking populations from South Africa (e.g. amaXhosa) as well as the admixed Western Cape SAC populations (de Wit *et al*. 2010), supporting a hypothesis of distinct *southern-specific* KhoeSan ancestry (Figure S1, S2) shared between indigenous and admixed groups.

**Figure 2:**
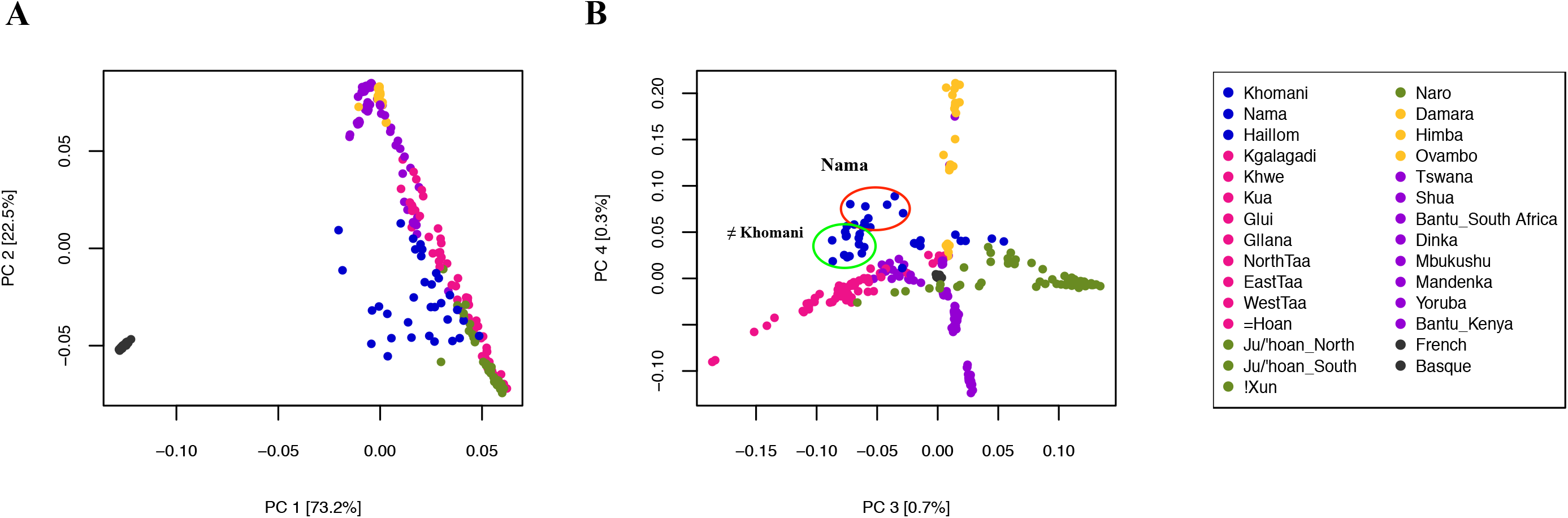
Clustering of KhoeSan populations and fine-scale population structure between the Nama and ≠Khomani San. A PCA of the Affymetrix Human Origins dataset depicts the clustering of unrelated individuals based on the variation seen in the dataset. The red and green circles indicate the fine-scale separation of the Nama and ≠Khomani populations.

Mitochondrial data support this concept of a *southern*-specific KhoeSan ancestry (Schlebusch *et al*. 2013; Barbieri *et al*. 2013). Both mtDNA haplogroups L0d and L0k are at high frequency in northern KhoeSan populations (Behar *et al*. 2008), but L0k is absent in our sample of the Nama [*n*=31] and there is only one ≠Khomani individual [*n*=64] with L0k (1.56%) **(Table 1).** L0d dominates both the Nama and ≠Khomani (84% and 91% respectively), with L0d2a especially common in both. L0d2a, inferred to have originated in southern Africa, was also previously found at high frequencies in the Karretjie people further south in the central Karoo of South Africa as well as the SAC population in the Western Cape (Quintana-Murci *et al*. 2010; Schlebusch *et al*. 2013). L0d2b is also common in the Nama (16%) (Schlebusch *et al*. 2013).

**Table 1.**
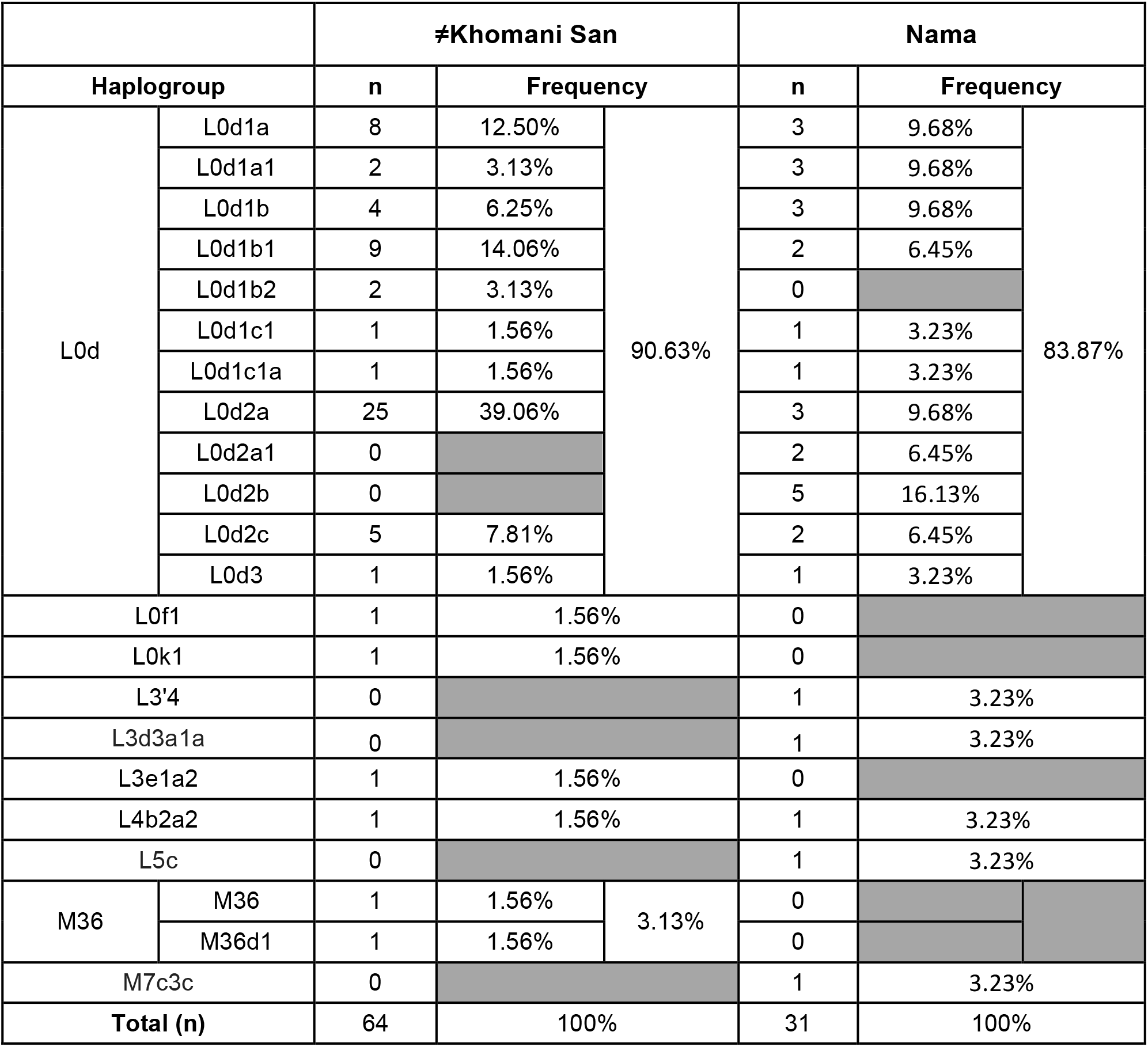
mtDNA haplogroup frequencies in the, ≠Khomani San and Nama popula:ons. Numbers of individuals is denoted by n.

### Reduced Population Structure Between the Nama and ≠Khomani

The ≠Khomani San are a N|u-speaking (!Ui classified language) hunter-gatherer population that inhabit the southern Kalahari Desert in South Africa, bordering on Botswana and Namibia. The Nama, a caprid pastoralist population, live in the Richtersveld along the northwestern coast of South Africa and up into Namibia. The ancestral geographic origin of the Nama has been widely contested over a number of years (Nurse and Jenkins 1977; Barnard 1992b; Boonzaier 1996), but a leading hypothesis suggests that they originated further north in Botswana/Zambia and migrated into South Africa and Namibia approximately 2,000 years ago (Nurse and Jenkins 1977; Barnard 1992b; Boonzaier 1996; Pickrell *et al*. 2012). The Nama and N|u languages are in distinct, separate Khoisan language families (Khoe and Tuu [!Ui-Taa], respectively) and these groups historically utilized different subsistence strategies. For this reason, we hypothesized that there would be strong population structure between the two populations.

Our global ancestry results, inferred from ADMIXTURE, show minimal population structure between the Nama and ≠Khomani San in terms of their southern KhoeSan ancestry. The ≠Khomani share ∽10% of their ancestry with the Botswana KhoeSan populations (Figure S2, S3), consistent with their proximity to the central Botswana populations (the Shua, Kua, G|ui and G||ana). Principal components analysis reveals a degree of fine-scale population structure between the Nama and ≠Khomani, with each population forming its own distinct cluster at PC4, partly due to the increase in Damara ancestry in the Nama **(Figure 2b,** Figure S1), but the groups are clearly proximal. This increase in Damara ancestry (as depicted from k=9 in all modes of Figure S1) is likely due to the enslavement of the Damara people by the Nama over multiple generations.

### Recent Patterns of Admixture in South Africa

Two Bantu-speaking, spatially distinct ancestries are present in southern Africa. The first is rooted in the Ovambo and Himba in northwestern Namibia; the other reflects gene flow of Bantu-speaking ancestry from the east **(Figure 1).** We estimated the time intervals for admixture events into the southern KhoeSan via analysis of the distribution of local ancestry segments using RFMix (Maples *et al*. 2013) and TRACTs (Gravel 2012) for the ≠ Khomani OmniExpress dataset (*n*=59 unrelated individuals) **(Figure 3).** The highest likelihood model suggested that there were 3 gene flow events. Approximately 14 generations ago (∽443-473 years ago assuming a generation time of 30 years and accounting for the age of our sampled individuals), the ≠Khomani population received gene flow from a Bantu-speaking group, represented here by the Kenyan Luhya. Our results are consistent with Pickrell et al. (2012) who found that the southern Kalahari Taa-speakers were the last to interact with the expanding Bantu-speakers about 10-15 generations ago. Subsequently, this event was followed by admixture with Europeans between 6 and 7 generations ago (∽233-263 years ago), after the arrival of the Dutch in the Cape and the resulting migrations of “trekboers” (nomadic pastoralists of Dutch, French and German descent) from the Cape into the South African interior. Lastly, we find a recent pulse of primarily KhoeSan ancestry 4-5 generations ago (∽173-203 years ago). This event could be explained by gene flow into the ≠Khomani from another KhoeSan group, potentially as groups shifted local ranges in response to the expansion of European farmers in the Northern Cape, or other population movements in Namibia or Botswana.

**Figure 3:**
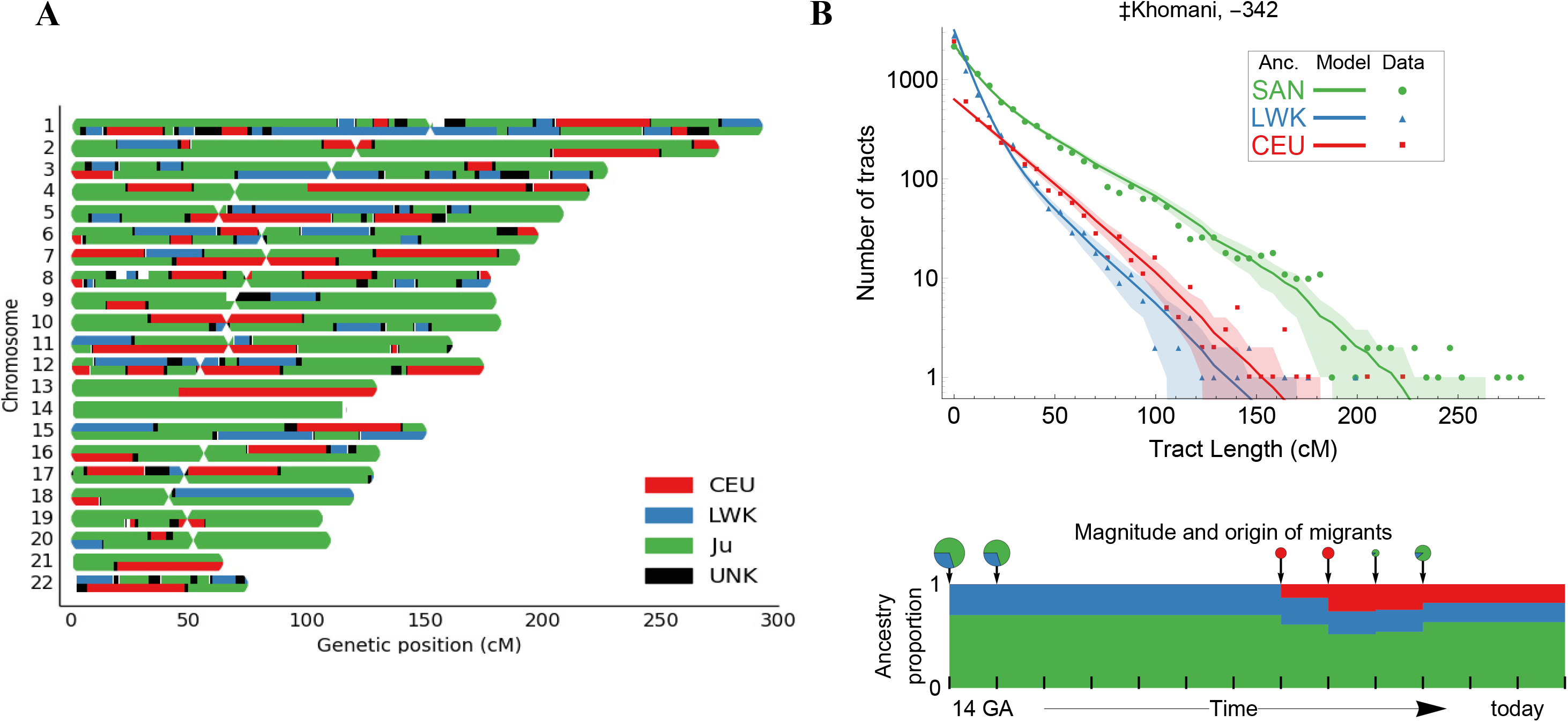
Demographic reconstruction of recent admixture in the ≠Khomani San. A) Local ancestry karyogram for a representative 3-way admixed ≠Khomani San individual using RFMix. B) We employed Markov models implemented in TRACTs to test multiple demographic models and assess the best fit to the observed data. The tract length distribution was used to estimate migration time (generations ago), volume of migrants, and ancestry proportions over time. Points show the observed distribution of ancestry tracts for each ancestry, solid lines show the best fit from the most likely model, and shaded areas show confidence intervals corresponding to ± 1 standard deviation.

We also considered the impact of recent immigration into indigenous South Africans, derived from non-African source populations. The South African Coloured (SAC) populations are a five way admixed population, deriving ancestries from Europe, east African, KhoeSan, and Asian populations (de Wit *et al*. 2010). This unique, admixed ethnic population was founded as a result of migrations and settlers moving into South Africa starting in the 6^th^ Century with the arrival of Bantu-speaking populations followed by the Dutch who settled on the southern tip of South Africa by the 17^th^ century. However, within this ethnic designation, strong differences in ancestry and admixture proportions are observed between different districts within Cape Town, the Eastern Cape and the Northern Cape Provinces. South African Coloured individuals from the Northern Cape, where historically there was a greater concentration of European settlement (Theal 1887), have higher European ancestry. The SAC individuals from the Eastern Cape, which is the homeland of the Bantu-speaking Xhosa populations, have relatively more ancestry from Bantu-speaking populations (Figure S2). The “ColouredD6” population is from an area in Cape Town called District 6. Historically, this was a district where the slaves and political exiles from present day Indonesia resided, who were primarily from Madagascar and India based on written documentation (Plessis 1947). The SAC D6 population consequently has a noticeable increase in southeastern Asian ancestry represented by the Pathan population in our dataset (Figure S2).

This South / East Asian ancestry is not confined to the SAC population, as attested by the presence of the M36 mitochondrial haplogroup. The M36 mitochondrial haplogroup (South Indian/Dravidian in origin) is present in two out of 64 ≠Khomani San matrilineages, **(Table 1).** The presence of M36 is likely derived from slaves of South Asian origin who escaped from Cape Town or the surrounding farms and dispersed into the northwestern region of South Africa. In addition, we observe one M7c3c lineage in the Nama **(Table 1),** which traces back to southeastern Asia but has been implicated in the Austronesian expansion of Polynesian speakers into Oceania (Kayser 2010; Delfin *et al*. 2012) and Madagascar (Poetsch *et al*. 2013). The importation of Malagasy slaves to Cape Town may best explain the observation of M7c3c in the Nama.

## Discussion

The African KhoeSan are distinguished by their phenotype, genetic divergence, click languages and hunter-gatherer subsistence strategy; classifications of the many ethnic groups have primarily relied on linguistic classification or subsistence strategy. Here, we generate additional genome-wide data from 3 South African populations and explore patterns of fine-scale population structure among 21 southern African groups. We find that complex geographic or “ecological” information is likely a better explanatory variable for genetic ancestry than language or subsistence. We identify 5 primary ancestries in southern Africans, each localized to a specific geographic region **(Figure 1).** In particular, we examined the “circum-Kalahari” which appears as a ring around the Kalahari Desert and accounts for the primary ancestry of the Nama, representative of the Khoekhoe-speaking pastoralists.

The practice of sheep, goat and cattle pastoralism in Africa is widespread. Within KhoeSan populations, pastoralist communities are limited to the Khoekhoe-speaking populations. Prior hypotheses proposed that the Khoe-speaking pastoralists were derived from a population originating outside of southern Africa. However, more recent genetic work supports a model of autochthonous Khoe ancestry influenced by either demic or cultural diffusion of pastoralism from East Africa ∽2,500 years ago (Pleurdeau *et al*. 2012; Pickrell *et al*. 2014). The presence of lactase persistence alleles in southern Africa was explained by Breton et al. (Breton *et al*. 2014), under two possible migratory scenarios. The first is a migration event from East Africa into southwestern Africa by the ancestors of the Khoe (Breton *et al*. 2014). The second is that there was contact between East African herders and Khoe populations in south-central Africa, with subsequent migration into southwest Namibia. This is also supported by Y-chromosomal analysis that indicates a direct interaction between eastern African populations and southern African populations approximately 2,000 years ago (Henn *et al*. 2008).

Our samples from the Khoekhoe-speaking Nama pastoralists demonstrate shared ancestry with other far southern non-pastoralist KhoeSan such as the ≠Khomani San and the Karretjie (see also Schlebusch *et al*. 2011). mtDNA also suggests that the Nama display a haplogroup frequency distribution more similar to KhoeSan south of the Kalahari than to any other population in south-central Africa. Our results indicate that the majority of the Nama ancestry has likely been present in far southern Africa for longer than previously assumed, rather than a recent migration from further north in Botswana where other Khoe-speakers live. The only other Khoekhoe-speaking population in our dataset is the Hai||om who share approximately 50% of the circum-Kalahari ancestry with the Nama and ≠Khomani, but are foragers rather than pastoralists. We conclude that Khoekhoe-speaking populations share a circum-Kalahari genetic ancestry with a variety of other Khoe-speaking forager populations in addition to the !Xun and ≠Khomani (**Figures 1** and **2**) rather than one defined by specific language families. This ancestry is divergent from central and northern Kalahari ancestries, arguing *against* a major demic expansion of Khoekhoe pastoralists from northern Botswana into South Africa. Rather, in this region, cultural transfer likely played a more important role in the diffusion of pastoralism. This is a unusual case of cultural transmission. Other economic transitions have been shown to be largely driven by demic diffusion (Gignoux *et al*. 2011; Fort 2012; Skoglund *et al*. 2014; Lazaridis *et al*. 2014; Malmström *et al*. 2015). Of course, a demic expansion of the Khoekhoe within Namibia and South Africa may still be have occurred - but geneticists currently lack representative DNA samples from many of the now “Coloured” interior populations which may carry Khoekhoe ancestry.

The transfer of pastoralism from eastern to southern Africa was not purely cultural and we can infer the effect of recent gene flow. For example, both the E3b1-M293 Y-chromosome lineage and the presence of East African lactase persistence alleles indicate an eastern African signature found across KhoeSan populations (though not exclusive to contemporary pastoralists) (Henn *et al*. 2008; Ranciaro *et al*. 2014; Breton *et al*. 2014; Macholdt *et al*. 2014). We also report here the presence of mitochondrial L4b2 that supports limited gene flow from eastern Africa. L4b2, formerly known as L3g or L4g, is a mtDNA haplogroup historically found at a high frequency in eastern Africa, in addition to the Arabian Peninsula. L4b2 is at high frequency specifically in click-speaking populations such as the Hadza and Sandawe in Tanzania (sometimes described as ‘Khoisan-speaking’) (Knight *et al*. 2003). Nearly 60% of the Hadza population and 48% of Sandawe belong to L4b2 (Tishkoff *et al*. 2007). Even though both Tanzanian click-speaking groups and the southern African KhoeSan share some linguistic similarities and a hunter-gatherer lifestyle, they have been isolated from each other over the past *35ky* (Tishkoff *et al*. 2007). The L4b2a2 haplogroup is present at a low frequency in both the Nama and ≠Khomani San, observed in one matriline in each population **(Table 1).** L4b2 was also formerly reported in the SAC population (0.89%) (Quintana-Murci *et al*. 2010) but has not been discussed in the literature. We identified several additional southern L4b2 haplotypes from whole mtDNA genomes deposited in public databases (Behar *et al*. 2008; Barbieri *et al*. 2013) and analyzed these samples together with all L4b2 individuals available in NCBI. Median-joining phylogenetic network analysis of the mtDNA haplogroup, L4b2, supports the hypothesis that there was gene flow from eastern Africans to southern African KhoeSan groups. As shown in Figure 4 (and in more detail in Figure S4), southern African individuals branch off from eastern African populations in this network (Salas *et al*. 2002; Tishkoff *et al*. 2007; Gonder *et al*. 2007). The mitochondrial network suggests a recent migratory scenario (estimated to be < 5,000 years before present), though the source of this gene flow, whether from eastern African click-speaking groups or others, remains, unclear (Pickrell *et al*. 2014).

**Figure 4:**
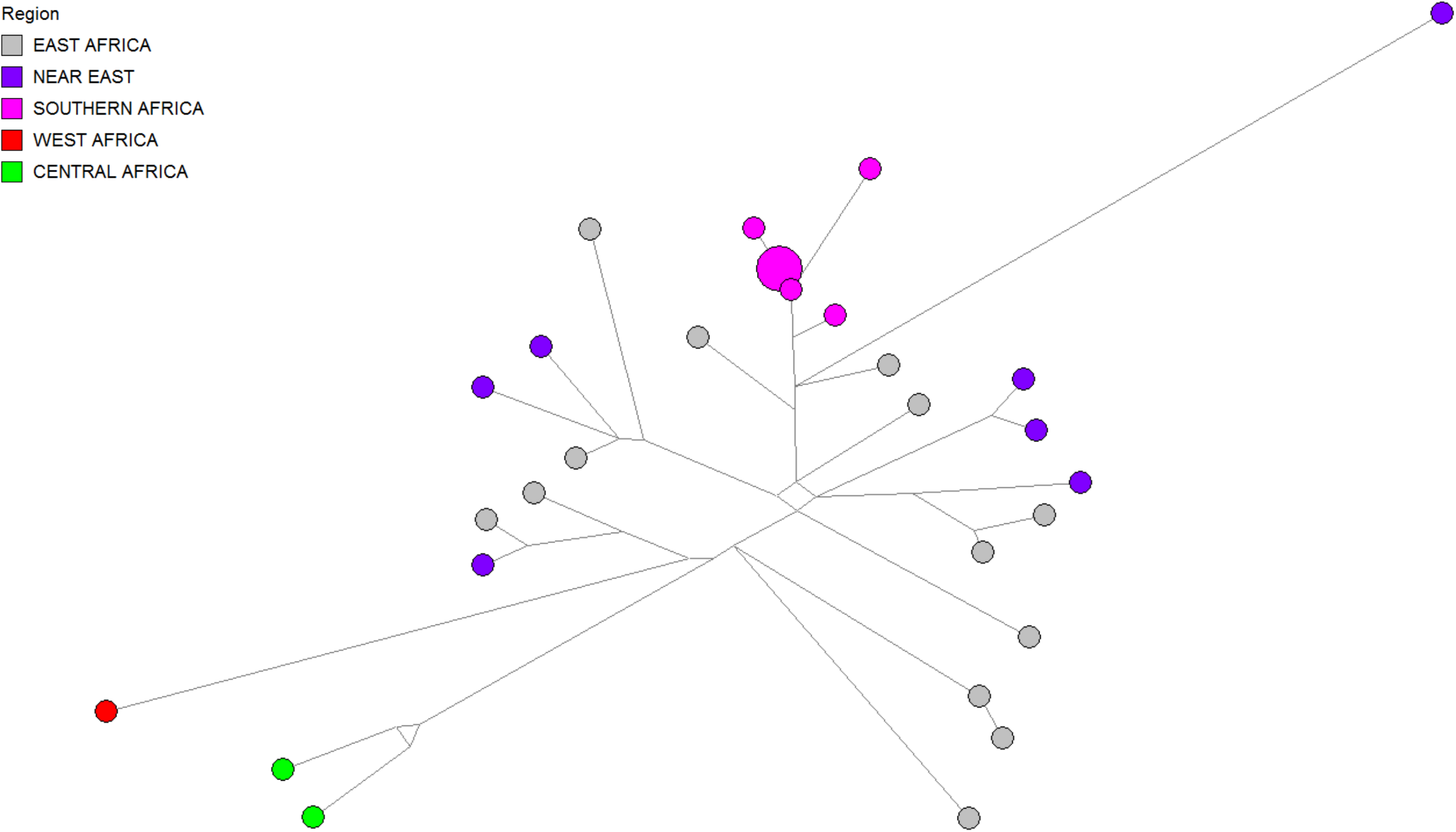
L4b2 mtDNA haplogroup network. ≠Khomani and Nama individuals, indicated in pink as “Southern Africa”, were analyzed together with publically available L4b2 mtDNA genomes from NCBI (as outlined in the Supplementary Methods). All individuals were assigned mtDNA haplogroups using *haplogrep* and the haplotypes were ploLed using Network Publisher.

## Conclusion

Analysis of 21 southern African populations reveals that fine-scale population structure corresponds with ecological rather than linguistic or subsistence categories. The Nama pastoralists are autochthonous to far southwestern Africa rather than representing a recent movement from further north. We find that the KhoeSan ancestry remains highly structured across the region and suggests that cultural diffusion likely played the key role in adoption of pastoralism.

## Materials and Methods

**Sample collection and ethical approval:** DNA samples from the Nama, ≠Khomani San and South African Coloured populations were collected with written informed consent and approval of the Human Research Ethics Committee of Stellenbosch University (N11/07/210), South Africa, and Stanford University (Protocol 13829), USA. Community level results were returned to the communities in 2015 prior to publication. A contract for this project was approved by WIMSA (ongoing).

### Autosomal data and genotyping platforms

A) ∽565 000 SNP Affymetrix Human Origins SNP array dataset from Pickrell et al. (Pickrell *et al*. 2012), Lazaridis et al. (Lazaridis *et al*. 2014) and additional ≠Khomani San and Hadza individuals from our collections: 33 populations and 396 individuals. B) ∽320,000 SNP array dataset from the intersection of HGDP (Illumina 650Y) (Li *et al*. 2008), HapMap3 (joint Illumina Human 1M and Affymetrix SNP 6.0), Illumina OmniExpressPlus and OmniExpress SNP array platforms as well as the dataset from Petersen et al. (Petersen *et al*. 2013): 21 populations and 852 individuals.

### Population structure

ADMIXTURE (Alexander *et al*. 2009) was used to estimate the ancestry proportions in a model-based approach. Iterations through various *k* values are necessary. The *k* value is an estimate of the number of original ancestral populations. Cross-validation (CV) was performed by ADMIXTURE and these values were plotted to acquire the *k* value that was the most stable. Depiction of the Q matrix was performed in R. Ten iterations were performed for each *k* value with ten random seeds. Iterations were grouped according to admixture patterns to identify the major and minor modes by pong (Behr *et al*. 2015). These Q matrixes from ADMIXTURE, as well as longitude and latitude coordinates for each population were adjusted to the required format for use in an R script supplied by Prof. Ryan Raaum to generate the surface maps (Figure 1).

### Association between F_st_, geography and language

A Mantel test (F_st_ and geographic distance) and a partial Mantel test (F_st_ and language, accounting for geographic distance) was performed using the vegan package in R. Geographic distances (in kilometers) between populations were calculated using latitude and longitude values as tabulated in Figure S1. Weir and Cockerham genetic distances (F_st_) were calculated from allele frequencies estimated with vcftools (Danecek *et al*. 2011). A Jaccard phonemic distance matrix was used as formulated in Creanza et al. (Creanza *et al*. 2015). Populations included in the analysis were the Nama, ≠Khomani, EastTaa, WestTaa, Naro, G|ui, G||ana, Shua, Kua, !Xuun and Khwe.

### mtDNA Network

We utilized Network (ver. 4.6, copy righted by Fluxus Technology Ltd.), for a median-joining phylogenetic network analysis in order to produce Figures 4 and S4. Network Publisher (ver. 2.0.0.1, copy righted by Fluxus Technology Ltd.) was then used to draw the phylogenetic relationships among individuals.

## Supplemental Information

Supplemental Information includes Supplemental Data, Supplemental Experimental Procedures, 4 figures and 3 tables.

## Author Contributions

C.U, M.K. A.R.M, and D.B. performed analysis. C.G., M.M., A. R.M, C.U., B.M.H. collected DNA samples. A.B. and S.R. contributed novel analysis tools. P.V.H, M.M, E.H and B.M.H. conceived of the study. C.U., C.R.G, M.M., E.H and B.M.H. wrote the manuscript in collaborations with all co-authors. All authors read and approved of the manuscript.

## Acknowledgements

Funding was provided by a Stanford University CDEHA seed grant to B.M.H. (NIH, NIA P30 AG017253-12) as well as a Stanford University Computation, Evolutionary, and Human Genomics (CEHG) Trainee Research Grant to A.R.M. C.U. was funded by the National Research Foundation of South Africa. C.R.G was funded by T32HG. We thank Jeffrey Kidd for assisting with genotyping of samples. We thank David Poznik for providing off-target mtDNA reads from a separate next-generation sequencing experiment. We thank Aaron Behr and Sohini Ramachandran for pre-publication use of *pong* and Meng Lin for help with analyses. We thank Carlos Bustamante for his encouragement and support of this project. We thank Marcus Feldman for a close reading of our manuscript. We thank Julie Granka, Justin Myrick and Cedric Werely for assistance with saliva sample collection and Ben Viljoen for DNA extractions. Guidance from Ryan Raaum with regards to formulating the surface plots is appreciated. We would like to thank the Working Group of Indigenous Minorities in Southern Africa (WIMSA) and the South African San Institute (SASI) for their encouragement and advice. Finally, we thank Richard Jacobs, Willem de Klerk, Hendrik Kaiman, and the communities in which we have sampled; without their support, this study would not have been possible.

